# PPARγ brakes cytosolic DNA-induced macrophage inflammation by super-enhancer metabolic reprogramming

**DOI:** 10.64898/2026.10.02.756223

**Authors:** Xinyi Fan, Kai Gao, Ping Meng, Jingmei Xu, Gongrui Zhang, Yumei Yuan, Qifeng Gao, Zhuoyuan Ma, Fan Yang, Yongqiang Zhang, Yan Wang, Shiqiang Zhang

## Abstract

Cytoplasmic DNA sensing triggers intense innate immune responses that require tight regulation. PPARγ is a master metabolic regulator in macrophages, but its role in cytoplasmic DNA immunity remains unclear. Using poly(dA:dT) to mimic cytoplasmic DNA stimulation, we combined transcriptomics, epigenomics (PPARγ/H3K27ac ChIP-seq), and functional assays. Poly(dA:dT) induced a cytokine storm with MAPK activation and apoptosis/necrosis. PPARγ activation by rosiglitazone (RGZ) antagonized this response, reducing cell death and transcriptionally inhibiting MAPK genes (e.g., DUSP4/10). Mechanistically, PPARγ bound promoters/enhancers via PPRE motifs and distal intergenic regions, remodeling H3K27ac (reduced at inflammatory loci, increased at metabolic genes). RGZ induced de novo super-enhancers coupling PPAR metabolic programs (e.g., CPT2) with immune suppression by targeting TLR4, CEBPB, and PIM3. Thus, PPARγ acts as a brake in cytoplasmic DNA immunity, integrating metabolic reprogramming and inflammation suppression through a multi-level genomic/epigenetic program. This study highlights PPARγ as a key immune-metabolic integrator and therapeutic target for diseases driven by excessive cytoplasmic DNA sensing.

## Introduction

The innate immune system senses cytoplasmic DNA through pattern recognition receptors, triggering intense inflammatory responses including type I interferon responses and activation of MAPK/NF-κB signaling pathways(Gulen et al, 2023), which are essential to fight infection but are also finely regulated to prevent self-injury(Abe & Barber, 2014). Metabolic states are deeply intertwined with immune activation, and expression of nuclear receptor PPARγ as a central regulator of metabolic homeostasis in macrophages suggests that it may be a key molecule linking these two systems(Hernandez-Quiles et al, 2021). Although PPARγ agonists have been widely reported to inhibit inflammation triggered by lipopolysaccharides(Paukkeri et al, 2013), the mechanism of action in cytoplasmic DNA-specific immune responses remains unknown. In particular, is this inhibition a downstream effect of metabolic changes or is PPARγ directed at transcriptional reprogramming of inflammatory signaling pathways(Yu et al, 2023; Zhang & Jagannath, 2025)? Furthermore, whether PPARγ, as a transcription factor, exercises this regulation by occupying specific genomic elements (such as enhancers) or even remodeling higher-order chromatin structure remains unexplored (Abdel-Rasol & El-Sayed, 2025; Czimmerer & Nagy, 2023).

The aim of this study is to systematically analyze the regulatory role and mechanism of PPARγ in cytoplasmic DNA-triggered innate immunity. Using poly(dA:dT) to simulate cytoplasmic DNA stimulation, combined with transcriptomics, epigenomics and functional experiments, we found that PPARγ acts as a key brake signal after being specifically activated. It directly binds to and inhibits transcription of core inflammatory pathway genes such as MAPK. More importantly, we revealed the multi-level genomic regulatory blueprint for PPARγ to achieve this function: From direct intervention in conjunction with promoter regions, to widespread recruitment to remote classical enhancers to co-regulate gene networks, and finally through the construction of super enhancers to simultaneously strengthen its own metabolic program and suppress inflammatory signals. These findings establish a new paradigm for PPARγ as an immune-metabolic integrator, which uses a programmed, spatially multi-level epigenetic transcription program to inhibit excessive immune activation triggered by cytoplasmic DNA.

## Results

### Poly(dA:dT) stimulation elicits a strong multi-layered inflammatory response in macrophages and extensively activates MAPK signaling pathways

Poly(dA:dT), as a classic agonist of the cytoplasmic DNA sensing pathway, has been reported to induce the interferon response and activate the MAPK pathway in various immune cells(Neamatallah et al, 2019; Tubita et al, 2025). To ensure that this pathway is equally effective in our WSL-PPARγ macrophage model and to provide a reliable experimental foundation for further investigation of PPARγ’s regulatory role, we performed RNA-seq analysis of stimulated macrophages. Poly(dA:dT) stimulation triggered extensive transcriptional reprogramming, resulting in 324 upregulated genes and 10 downregulated genes (adjusted p-value ≤ 0.05; |log2FC| ≥ 1)**(Fig. 1A)**. Further analysis revealed that the immune response was multi-layered and high-intensity. First, cluster analysis revealed that the type I interferon response pathway was strongly activated, and a number of interferon-stimulated genes (ISGs) including ISG15, OAS2, and MX2 were significantly upregulated **(Fig. 1B)**, marking a classical antiviral response state. Second, there was a full-scale explosion of pro-inflammatory cytokines and chemokines expression **(Fig. 1C, D)**. KEGG pathway enrichment analysis was performed to systematically unravel the regulatory network of this innate immune response. The results showed that differentially expressed genes were significantly enriched in Toll-like receptor signaling, NF-κB signaling, JAK-STAT signaling, and MAPK signaling **(Fig. 1E)**. The aforementioned results confirm that poly(dA:dT) effectively activates key inflammatory pathways such as MAPK/NF-κB in our macrophage model, providing an experimental foundation for further investigating PPARγ’s regulatory role in this pathway. Among them, MAPK signaling pathway is a key hub for regulating cell proliferation, stress, inflammation, and death, and its enrichment attracted our attention(Moens et al, 2013). To further verify the activation status of MAPK pathway, we focused on its three main branches-ERK, JNK and p38. Heat map analysis clearly showed that 14 key genes in ERK, JNK and p38 pathway was significantly upregulated after poly(dA:dT) stimulation **(Fig. 1F-H)**. Collectively, these results demonstrate that poly(dA:dT) elicits a strong and multifaceted inflammatory response in macrophages, in which MAPK signaling has been identified as a key activation hub.

**Figure 1.**
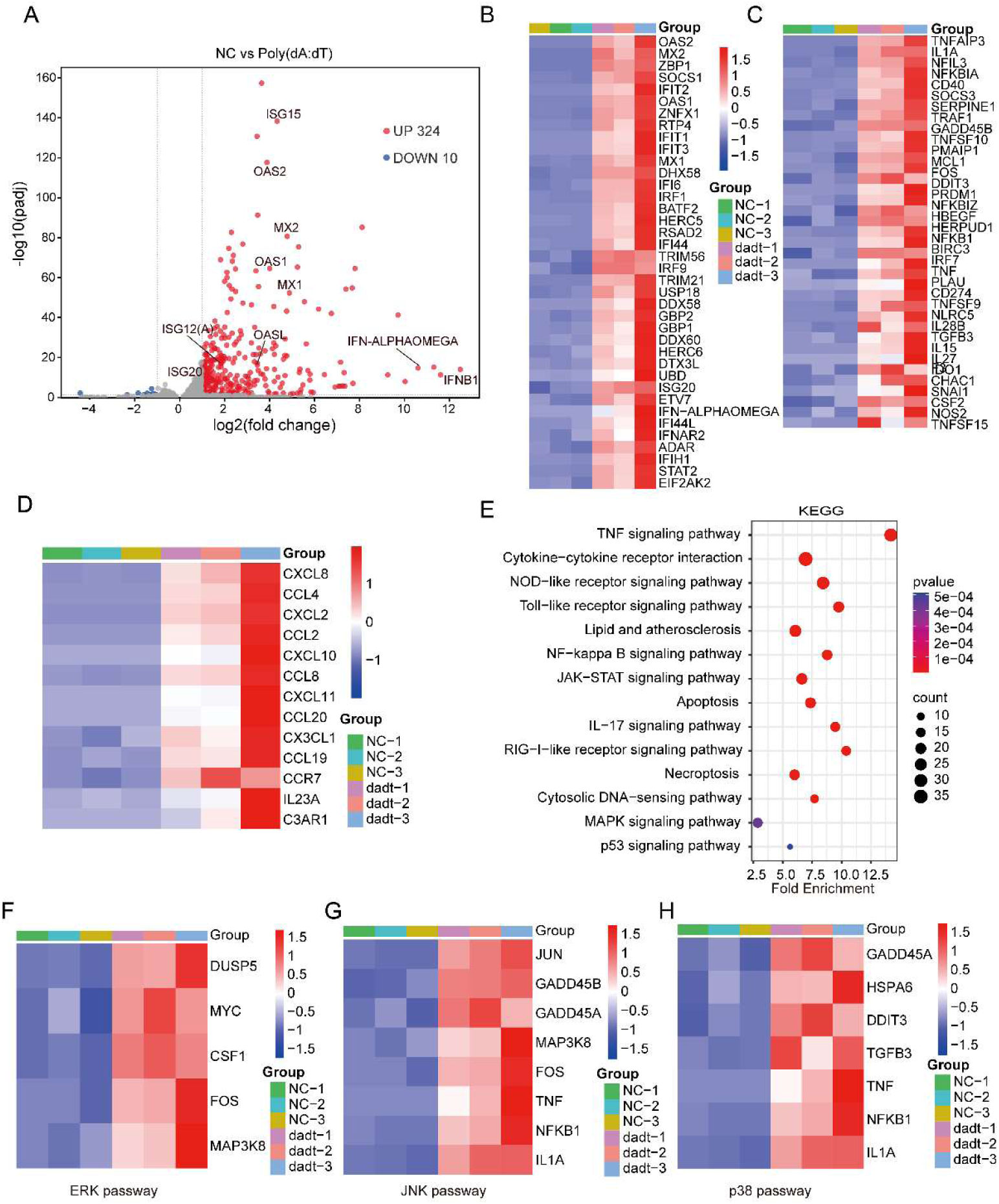
Transcriptomic changes and signaling pathway analysis induced by poly(dA:dT) stimulation. **(A)** Volcano plot of differentially expressed genes in macrophages after poly(dA: dT) stimulation. Differentially expressed genes were identified using the following criteria: padjust ≤ 0. 05; log2FC ≥ 1. Upregulated genes are indicated by red dots, and downregulated genes by blue dots.**(B)** Heatmap showing the expression patterns of interferons (IFNs) and their downstream classical interferon-stimulated genes (ISGs) induced by poly(dA:dT) stimulation.**(C)** Heatmap showing the expression patterns of cytokines and their downstream responsive genes induced by poly(dA:dT) stimulation.**(D)** Heatmap showing the expression patterns of chemokines and their related genes induced by poly(d A:dT) stimulation.**(E)** KEGG pathway enrichment analysis of differentially expressed genes after poly(d A:dT) stimulation. The top significantly enriched signaling pathways are shown. **(F)** Heatmap of ERK pathway-related gene expression. **(G)** Heatmap of JNK pathway-related gene expression. **(H)** Heatmap of p38 pathway-related gene expression. The RNA-seq experiment included three biological replicates, each with three technical replicates.

### PPARγ activation antagonizes inflammation and exerts cytoprotective effects through transcriptional inhibition of MAPK signaling pathway

Based on the intense inflammation induced by poly(dA:dT) and extensive activation of MAPK pathway, we hypothesized that there is an endogenous brake feedback regulatory mechanism in cells. PPARγ, as an important metabolic and immunoregulatory nuclear receptor, is a potential candidate factor(Yu et al., 2023). To elucidate its role, we first treated macrophages with PPARγ specific agonist rosiglitazone (RGZ) and antagonist T0070907 (T007), and evaluated the specific effect of PPARγ functional state on transcriptome by RNA-seq. Pharmacological intervention effectively and specifically altered PPARγ transcriptional activity. RGZ treatment elicited extensive gene expression changes (450 genes up-regulated and 150 genes down-regulated) compared to DMSO control, whereas T007 treatment induced relatively limited changes **(Fig. 2A-C)**. Direct comparison between RGZ and T007 groups confirmed the effectiveness of the intervention, including the strongly induced classical PPARγ target gene FABP4 **(Fig. 2C)**. KEGG enrichment analysis revealed that genes regulated by RGZ and T007 were co-enriched in PPAR signaling pathway and fatty acid metabolism pathway, consistent with their core functions **(Fig. 2D, E)**. It is worth noting that MAPK signaling pathway was also significantly enriched **(Fig. 2D)**, suggesting that PPARγ may be involved in regulating this pathway.

**Figure 2.**
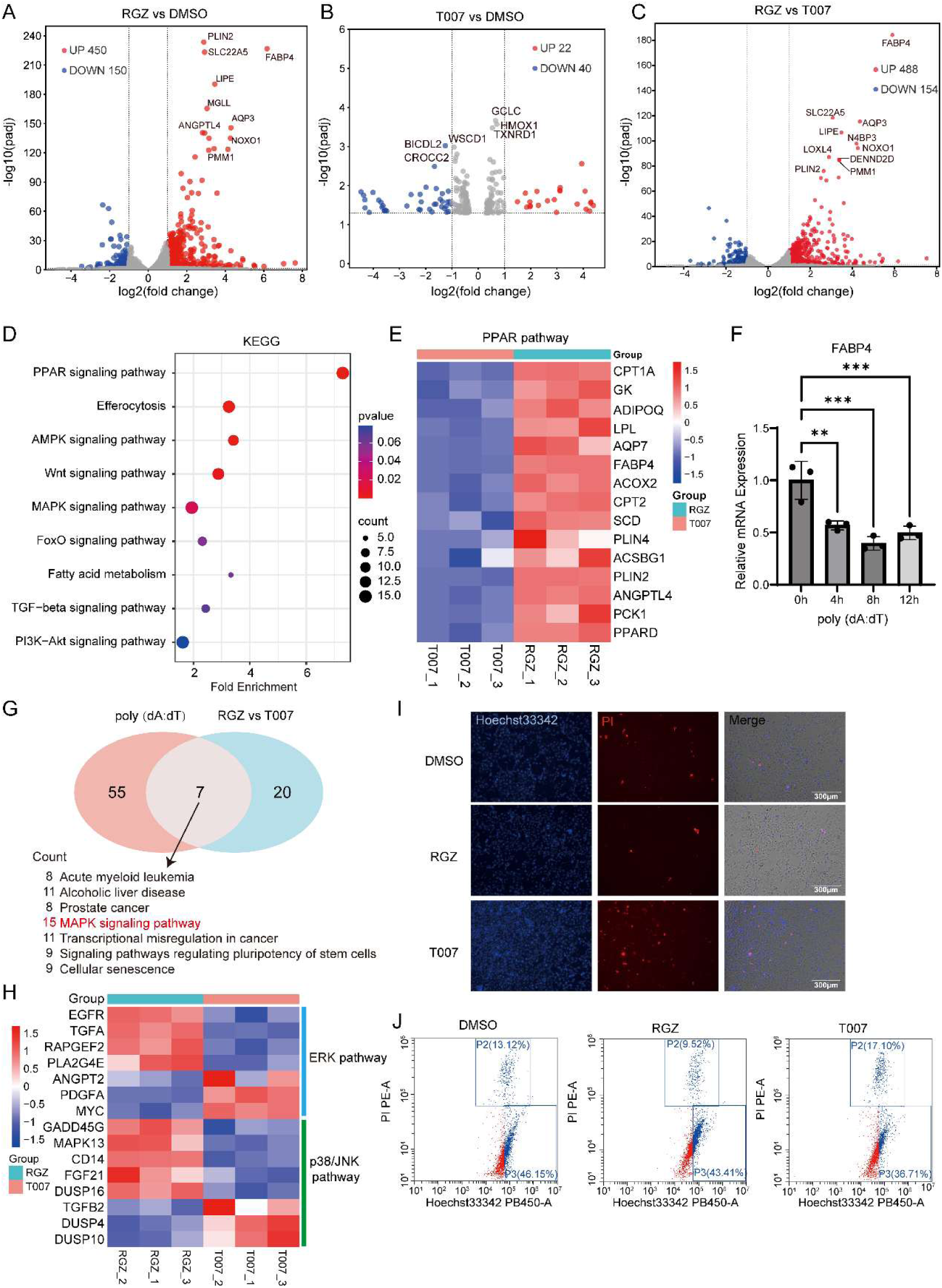
Effects of PPARG activation and inhibition on gene expression, signaling pathways, and cellular phenotypes. **(A)** Volcano plot of differentially expressed genes in poly(dA:d T)-stimulated macrophages treated with the PPARγ agonist rosiglitazone (RGZ) (padjust ≤ 0. 05; |log2 FC| ≥ 1). **(B)** Volcano plot of differentially expressed genes in poly(d A:dT)-stimulated macrophages treated with the PPARγ inhibitor T0070907 (T007). **(C)** Volcano plot of differentially expressed genes after RGZ and T007 treatment. **(D)** KEGG pathway enrichment analysis of differentially expressed genes after RGZ/T007 treatment. **(E)** Heatmap of PPAR signaling pathway-related gene expression under different treatments. **(F)** Expression changes of the PPARG target gene FABP 4 after poly(dA:dT) stimulation. **(G)** Venn diagram showing the intersection of KEGG pathways enriched under three conditions: poly(dA:dT) stimulation, RGZ-mediated PPARG activation, and T007-mediated PPARG inhibition. **(H)** Heatmap of the effects of RGZ/ T007 treatment on MAPK signaling pathway gene expression. **(I)** Apoptosis and necrosis assessed by Hoechst 33342/propidium iodide (PI) staining in poly(dA:dT)-stimulated macrophages treated with RGZ or T007. Scale bar: 300 μm. **(J)** Flow cytometric quantification of cell death. Annexin V-FITC/PI double staining was used to detect the effects of RGZ or T007 treatment on apoptosis and necroptosis after poly(dA:dT) stimulation. The RNA-seq experiment included three biological replicates, each with three technical replicates. Hoechst/PI staining and flow cytometry experiments were independently repeated three times.

We then explored the functional association between PPARγ regulation and poly(dA:dT) inflammatory stimulation. Interestingly, poly(dA:dT) stimulation itself inhibits FABP4 expression **(Fig. 2F)**, suggesting that inflammatory signaling may negatively regulate PPARγ basal activity. Venn diagram analysis revealed that MAPK signaling is the pathway with the largest number of overlapping target genes by systematically comparing pathways enriched by poly(dA:dT) stimulation, RGZ activation, and T007 inhibition **(Fig. 2G)**. This finding strongly suggests that MAPK signaling is a key hub for PPARγ regulation of poly(dA:dT)-triggered inflammatory responses. To test this hypothesis, we directly analyzed the effect of PPARγ regulation on MAPK pathway gene expression. Heatmap results showed that RGZ activation of PPARγ significantly downregulated a series of key genes in multiple branches of MAPK pathway (ERK, p38/JNK), such as MYC, DUSP4, and DUSP10, to name a few, compared with T007 treatment **(Fig. 2H)**.

Finally, we assessed the functional consequences of this transcriptional inhibition. Activation of PPARγ by RGZ significantly reduced the ratio of apoptosis to necrosis in poly(dA:dT)-induced inflammatory settings, as demonstrated by Hoechst33342/PI staining **(Fig. 2I)** and quantitative flow cytometry analysis **(Fig. 2J)**, whereas T007 treatment had no such protective effect. These results link PPARγ activation, MAPK pathway transcriptional inhibition, and ultimate cell fate, suggesting that PPARγ plays a critical cytoprotective role in DNA-triggered inflammatory responses by inhibiting gene expression of MAPK signaling pathways.

### PPARγ as a transcription factor directly binds to genes that regulate the origin of immune inflammatory signaling networks

To elucidate the direct transcriptional regulatory mechanism of PPARG, we conducted ChIP-seq analysis. First, we analyzed the binding patterns of PPARγ near the transcription start site (TSS) (**Fig. 3A, B**). Compared with the input control group, IP samples treated with RGZ or T007 exhibited specific enrichment signals in the regions upstream and downstream of the TSS, confirming that PPARγ directly binds to gene regulatory regions as a transcription factor. By comparing the binding signals between the RGZ and T007 groups, differential volcano plots identified a significant number of binding sites that were markedly enhanced after RGZ treatment (**Fig. 3C**). Motif analysis revealed that these sites were highly enriched with typical PPAR response elements (PPREs) (**Fig. 3D**), verifying the specificity of the binding. Genomic distribution analysis indicated that the differentially bound peaks were predominantly located in the intron and intergenic (**Fig. 3F**). To elucidate the biological functions of these target genes, we conducted GO and KEGG enrichment analyses on the differential binding peaks in the promoter regions. The GO analysis (**Fig. 3G**) revealed that the target genes were significantly involved in core immune and inflammatory processes such as “positive regulation of viral transcription”, “NIK/NF-κB signaling pathway”, “JAK-STAT signaling pathway” and “positive regulation of protein serine/threonine kinase activity”. Additionally, genes related to “positive regulation of cell cycle arrest” and “histone deacetylase complexes” were significantly enriched. The KEGG analysis (**Fig. 3H**) further demonstrated that these genes were markedly enriched in innate immune pathways, including the “cytoplasmic DNA sensing pathway” and “JAK-STAT signaling pathway”. The aforementioned ChIP-seq results indicate that activated PPARγ occupies the origin of the regulatory network by directly binding to the promoter regions of numerous key genes involved in immune and infection responses. Its target genes were significantly enriched in signaling axes directly associated with poly(dA:dT)-mediated inflammatory activation, such as cytoplasmic DNA sensing and JAK-STAT pathways, as well as broader infection response and immune regulation pathways. In situ binding assays of transcription factors confirmed that PPARγ serves as an upstream regulatory hub, directly intervening in the core inflammatory signaling networks triggered by pattern recognition receptors.

**Figure 3.**
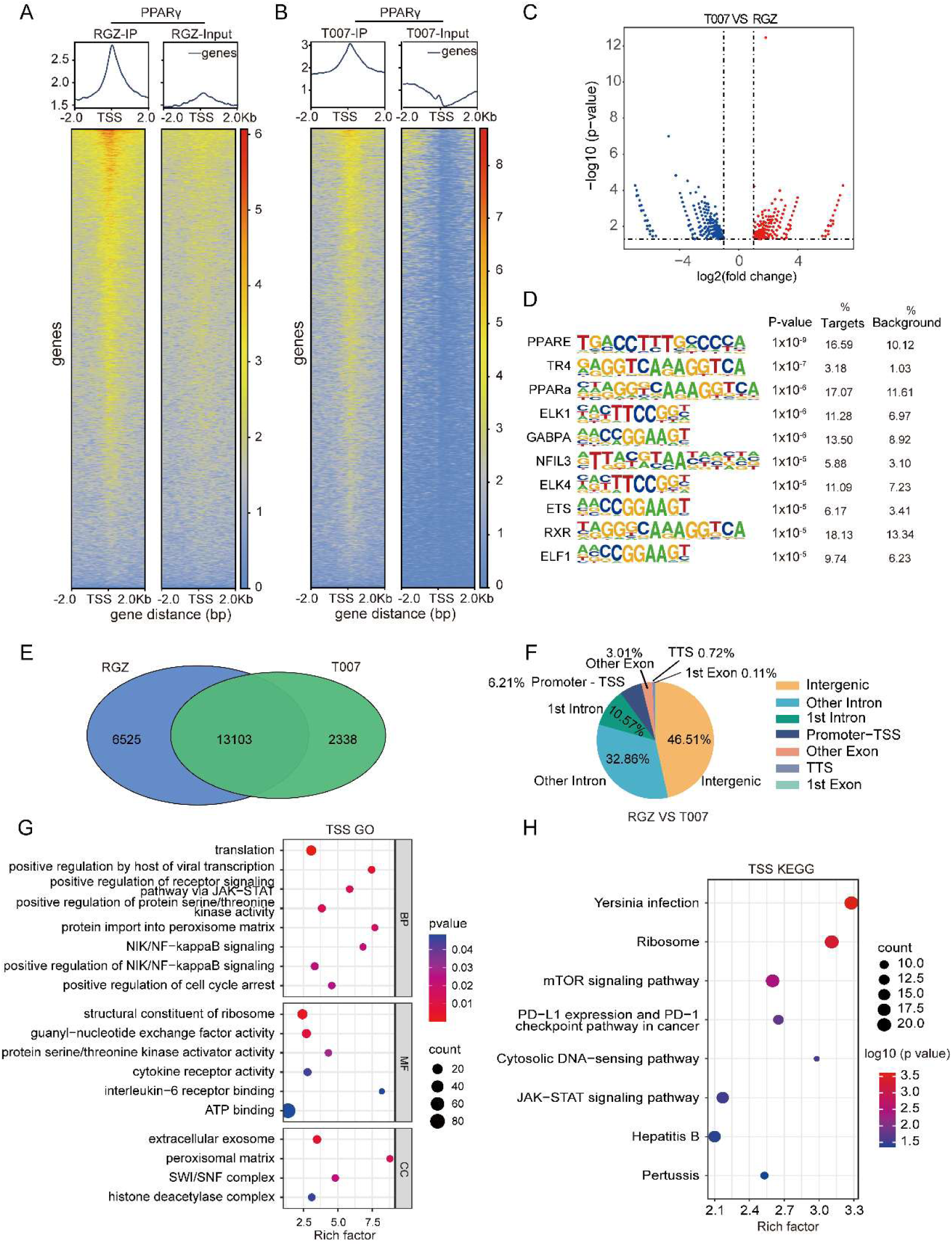
PPARγ ChIP-seq reveals its genomic binding landscape. **(A)** Genomic binding landscape of PPARγ under treatment with the PPARγ agonist rosiglitazone (RGZ). The upper panel shows the overall enrichment pattern of ChIP-seq signal (IP group) and Input control in regions upstream and downstream of the transcription start site (TSS); the lower panel shows a heatmap of normalized signal intensity for each peak around the TSS. **(B)** Genomic binding landscape of PPARγ under treatment with the PPARγ inhibitor T007. The presentation is the same as in **(A)**. **(C)** Volcano plot of differentially bound peaks between the RGZ and T007 treatment groups, showing differences in binding intensity and statistical significance. **(D)** Significantly enriched DNA sequence motifs among differentially bound peaks. **(E)** Venn diagram of differentially bound peaks between the RGZ and T007 treatment groups. The numbers of group-specific and shared PPARγ binding sites are shown. **(F)** Genomic distribution proportions of differentially bound peaks between the RGZ and T007 treatment groups. **(G)** Gene Ontology (GO) functional enrichment analysis of genes associated with differentially bound peaks. **(H)** KEGG pathway enrichment analysis of genes associated with differentially bound peaks. ChIP-seq experiments were performed using an anti-FLAG antibody to immunoprecipitate the PPARγ-FLAG fusion protein. Peak calling was performed using MACS2 software (q-value < 0.05). Functional enrichment analysis was based on the nearest genes associated with differentially bound peaks.

### PPARγ synergistically inhibits MAPK pathway and constructs protective transcription program by occupying distal classical enhancers

Based on the discovery that PPARγ binds directly to multiple upstream immune gene promoters, we further explored whether it coordinates the overall inflammatory program through broader genomic regulation. In-depth analysis of ChIP-seq data revealed 39.36% of PPARγ binding sites were located in the intergenic region after RGZ activation, significantly higher than in the T007 treatment group (28.69%) **(Fig. 4A, B)**. This suggests that PPARγ activation exerts its regulatory effects may span from proximal promoters to distal regulatory elements, such as classical enhancers.

**Figure 4.**
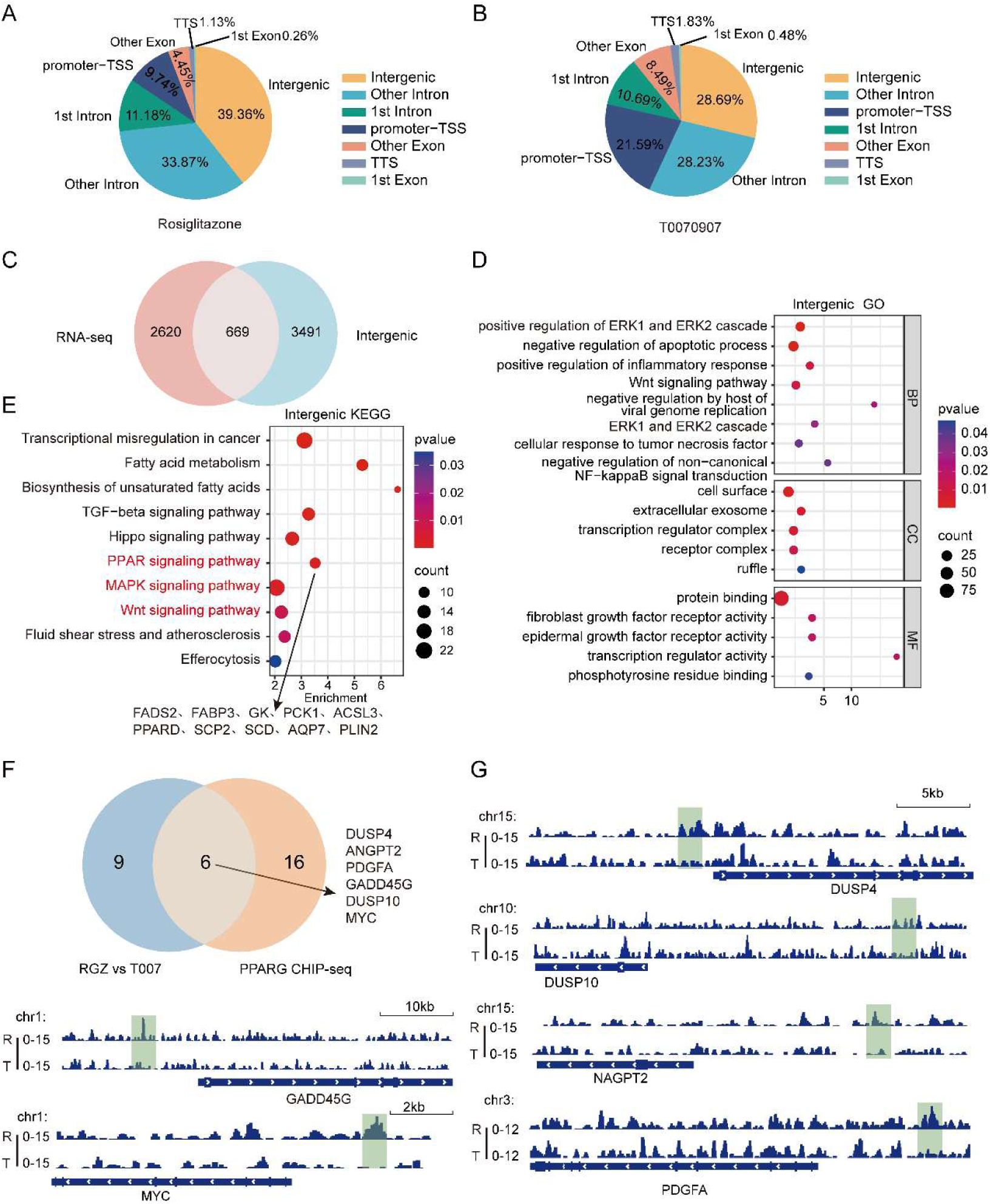
Integrated multi-omics analysis reveals that PPARγ affects PPAR and MAPK pathways by binding to non-coding regulatory regions. **(A)** Distribution proportions of identified PPARγ binding peaks across genomic functional regions after RGZ treatment. **(B)** Distribution proportions of identified PPARγ binding peaks across genomic functional regions after T007 treatment. **(C)** Venn diagram showing the intersection between genes associated with intergenic differentially bound PPARγ peaks and RNA-seq differentially expressed genes between the RGZ and T007 treatment groups. **(D)** GO enrichment analysis of the intersecting genes associated with intergenic peaks. **(E)** KEGG enrichment analysis of the intersecting genes associated with intergenic peaks. **(F)** Multi-omics intersection analysis of MAPK pathway-related genes. The Venn diagram shows the overlap of RNA-seq differentially expressed genes and PPARγ ChIP-seq target genes within the MAPK pathway. **(G)** Integrative Genomics Viewer (IGV) views of the six key intersecting genes from (F). ChIP-seq signals for PPARγ (FLAG) under RGZ and T007 treatments and gene annotations are simultaneously displayed, visually presenting the association of multi-omics data. All enrichment analyses were based on the hypergeometric distribution test, and a corrected P-value (padjust) ≤ 0. 05 was considered significant. Ch IP-seq target genes were defined as the nearest neighboring genes upstream and downstream of differentially bound peaks.

To verify the function of these distal binding sites, we correlated PPARγ differentially binding sites in the intergenic region with differentially expressed genes after RGZ treatment. Venn plots showed that 669 intergenic sites were significantly associated with genes with altered expression **(Fig. 4C)**. Functional analysis of these associated genes revealed that they were highly enriched in key processes such as “positive regulation of ERK1/ERK2 cascade”, “negative regulation of apoptosis” and “negative regulation of non-classical NF-κB signaling”**(Fig. 4D)**. KEGG pathway analysis further confirmed that these genes regulated by distal elements were significantly enriched in “MAPK signaling pathway”, “PPAR signaling pathway” and “Wnt signaling pathway” **(Fig. 4E)**.

In order to focus on the core regulatory mechanism of PPARγ on MAPK pathway, we intersected MAPK pathway genes associated with binding sites in intergenic region with MAPK pathway genes differentially expressed in RNA-seq, and identified six core regulatory genes, namely DUSP4, DUSP10, ANGPT2, PDGFA, MYC, and GADD45G **(Fig. 4F)**. It is worth noting that DUSP4 and DUSP10 are key negative feedback phosphatases of MAPK pathway(Neamatallah et al., 2019; Tubita et al., 2025). The finding that it is directly regulated by PPARγ has not been reported in the context of cytoplasmic DNA sensing. Through genome browser visualization, we clearly observed PPARγ binding specifically to distal regulatory region of these genes after RGZ treatment **(Fig. 4G)**.

The results in Supplementary Fig. S1 further complement and reinforce this multi-layered regulatory framework. Analysis of intronic PPARγ binding sites revealed that 790 intronic peaks were associated with differentially expressed genes (Supplementary Fig. S1A). GO analysis revealed that these genes were significantly enriched in key immune regulatory processes, such as “activation of innate immune response,” “inflammatory response,” and “negative regulation of ERK1 and ERK2 cascade” (Supplementary Fig. S1B). KEGG analysis again highlighted the central role of the “PPAR signaling pathway” (Supplementary Fig. S1C). Furthermore, IGV visualization clearly demonstrated PPARγ binding to the intergenic (Supplementary Fig. S1D) and intronic (Supplementary Fig. S1E) regions of key genes in the PPAR signaling pathway (e.g., PPARD, ACOX2), confirming at the genomic level the direct transcriptional regulation of its core pathway by PPARG.

In conclusion, our results reveal that PPARγ through promoters and putative enhancers (introns and intergenic regions) synergistic genome occupation, co-constructs a protective transcriptional program that suppresses inflammatory signaling (especially MAPK pathway) and apoptosis.

### PPARγ constructs multi-layered and stable transcriptional suppressor program by remodeling local H3K27ac modifications

To investigate the functional consequences of broad genomic binding of PPARγ and its potential impact on chromatin status, we performed ChIP-seq analysis of H3 K27 acetylation (H3K27ac), which is an active enhancer marker. The results showed that RGZ treatment induced significant changes in local H3K27ac modifications. Compared with T007 treatment, RGZ induced extensive changes in H3K27ac signal, and the differential enrichment peaks were also mainly located in intergenic regions, highly coinciding with activated PPARγ binding patterns **(Fig. 5A-C)**. These results suggest that PPARγ binding is directly related to changes in local H3K27ac modifications.

**Figure 5.**
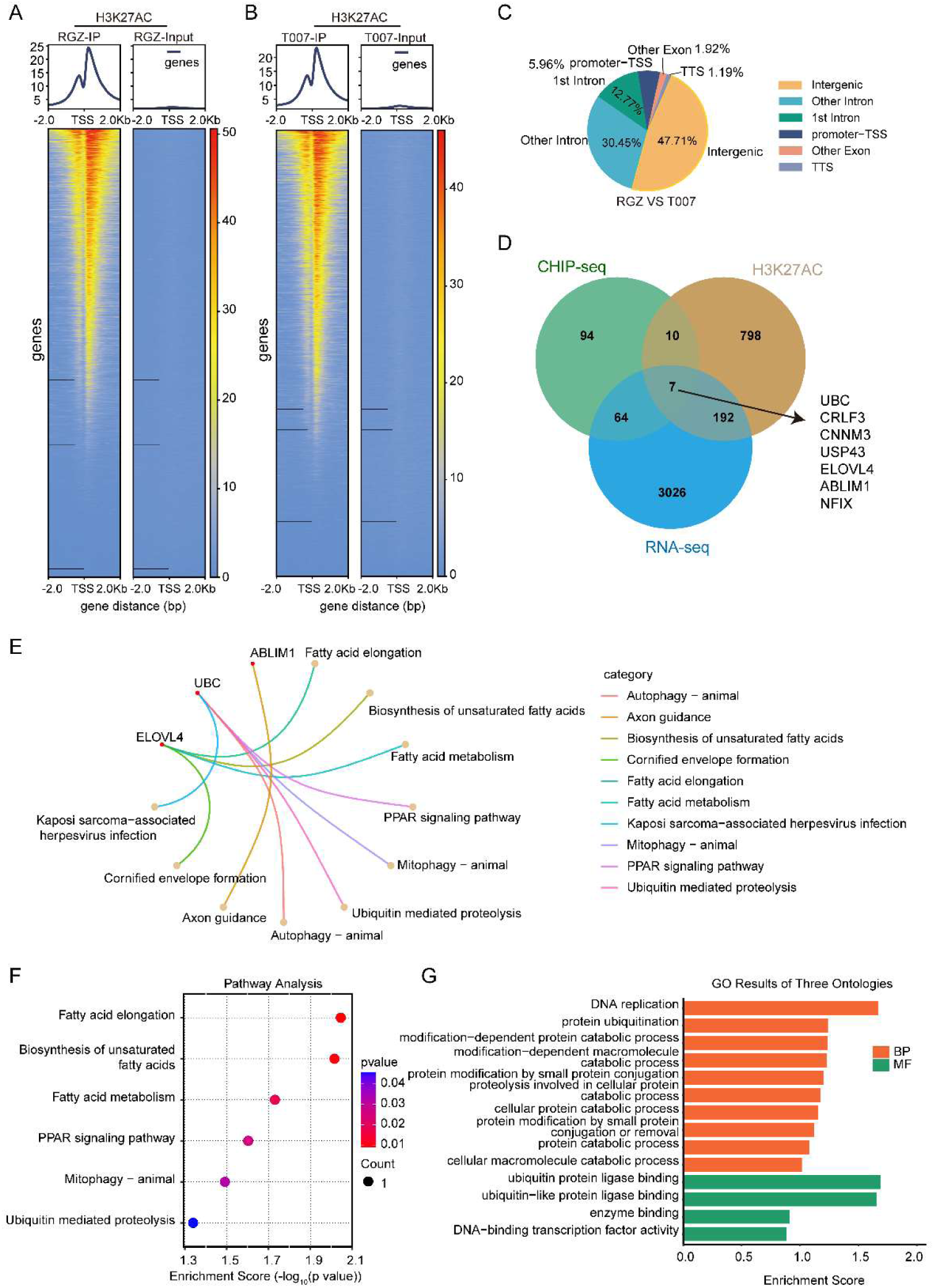
H3K 27ac modification landscape and multi-omics association analysis under PPARγ agonist and antagonist treatments. **(A)** Genomic enrichment landscape of the histone modification H3K27ac under treatment with the PPARγ agonist rosiglitazone (RGZ). The upper panel shows the overall enrichment pattern of H3K27ac ChIP-seq signal (IP) and Input control in regions upstream and downstream of the transcription start site (TSS); the lower panel shows a heatmap of normalized signal intensity for each peak around the TSS. **(B)** Genomic enrichment landscape of H3K27ac under treatment with the PPARγ inhibitor T007. The presentation is the same as in (A). **(C)** Distribution proportions of differentially bound H3K27ac peaks across genomic functional regions between the RGZ and T007 treatment groups. **(D)** Venn analysis of RNA-seq differentially expressed genes, PPARγ differentially bound genes (promoter-TSS regions), and H3K27ac differentially enriched genes (promoter-TSS regions), showing the set of co-regulated candidate target genes. **(E)** Distribution plot of KEGG pathway centrality (cent) for the intersecting genes obtained from the Venn analysis in (D). **(F)** Bubble plot of KEGG pathway enrichment analysis for the intersecting genes in (D). **(G)** Gene Ontology (GO) functional enrichment analysis of the intersecting genes in (D). A corrected P-value (padjust) ≤ 0. 05 was considered statistically significant for enrichment analysis.

To establish a complete causal chain of transcription factor binding-epigenetic remodeling-gene expression output, we integrated RNA-seq, PPARγ ChIP-seq and H3K27acChIP-seq data. Intersection analysis of promoter regions identified a chain of seven genes (UBC, CRLF3, CNNM3, USP43, ELOVL4, ABLIM1 and NFIX), whose promoters are simultaneously bound by PPARγ, undergo H3K27ac modification changes, and whose expression is regulated by RGZ **(Fig. 5D)**. Functional analysis revealed that these genes are mainly involved in “fatty acid metabolism”, “PPAR signaling pathway” and “ubiquitin-mediated proteolysis” **(Fig. 5E-G)**, revealing that PPARγ regulates core metabolic programs by directly binding and remodeling histone modifications in the promoter region.

In order to further elucidate the spatial regulatory network of PPARγ in the non-promoter regions and highlight the regulatory functions of intergenic and intronic regions, which have been rarely addressed in previous studies, we analyzed the binding function of PPARγ in intron and intergenic region (distal enhancer) **(Fig. 6A, B)**. In the intergenic region, the combination of genes associated with both inflammatory pathways (e.g. JAK-STAT) and metabolic reprogramming pathways (e.g. fatty acid metabolism) **(Fig. 6C, D)** reflects the integrated function of coordinated regulation of immunity and metabolism by remote enhancers. Finally, through focused multiomic integration of MAPK pathways, we identified key target genes such as DUSP4 and DUSP10 that simultaneously satisfy the three conditions of PPARγ binding, H3K27ac modification, and downregulation **(Fig. 6E)**. Genome visualization clearly shows the complete chain of regulatory evidence at these loci: PPARγ binding, concomitant H3K27ac signal attenuation, and corresponding transcriptional suppression **(Fig. 6F)**.

**Figure 6.**
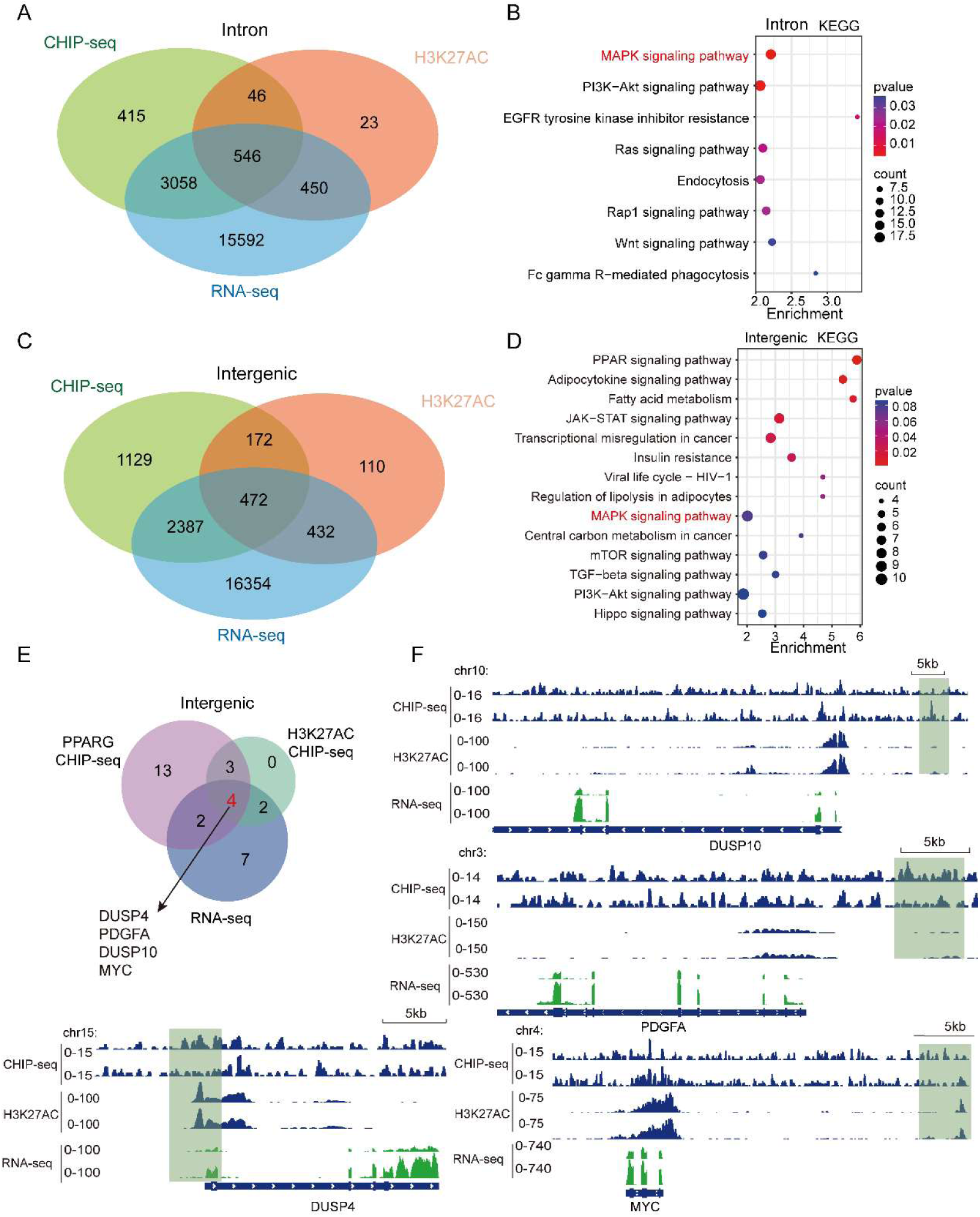
Analysis of PPARγ regulatory mechanisms based on genomic regions and multi-omics integration. **(A)** Venn analysis of intronic differentially bound peaks between RGZ and T007 treatments and RNA-seq differentially expressed genes, demonstrating the association between chromatin binding changes and transcriptional regulation within this region. **(B)** Bubble plot of KEGG pathway enrichment analysis for the intersecting genes obtained from the Venn analysis in (A). **(C)** Venn analysis of intergenic differentially bound peaks between RGZ and T007 treatments and RNA-seq differentially expressed genes. **(D)** Bubble plot of KEGG pathway enrichment analysis for the intersecting genes obtained from the Venn analysis in (C). **(E)** Venn diagram of intersecting genes among RNA-seq differentially expressed genes, PPARγ intergenic differentially bound genes, and H3K27ac intergenic differentially enriched genes within the MAPK signaling pathway, to screen for candidate regulatory target genes of this pathway. **(F)** IGV visualization of multi-omics signals for DUSP4, PDGFA, DUSP 10, and MYC from PPARγ and H3K27ac ChIP-seq and RNA-seq data. A corrected P-value (padjust) ≤ 0.05 was considered statistically significant for all enrichment analyses.

Our findings establish PPARγ binding can actively change the active state of local chromatin (e.g., reduce H3K27ac), which provides a stable chromatin basis for transcriptional suppression programs.

### PPARγ reshapes three-dimensional genome by constructing super enhancers, coupling metabolic programs with inflammation suppression

In order to explore the global regulatory capacity of PPARγ at a higher chromatin structure level, we performed a super enhancer analysis based on H3K27ac ChIP-seq data. Super enhancers are chromatin regulatory hubs formed by multiple classical enhancers, which are essential for maintaining cell characteristics and core gene expression(Jiang et al, 2021; Wang et al, 2023). The analysis found that 1191 super enhancers were identified in RGZ treatment group, significantly more than 919 in T007 treatment group **(Fig. 7A, B)**. Comparative analysis showed that a large number of unique super enhancers were produced under RGZ conditions **(Fig. 7C)**. This indicates that PPARγ activation can widely establish new higher-order chromatin regulatory hubs and achieve remodeling of genome three-dimensional structure.

**Figure 7.**
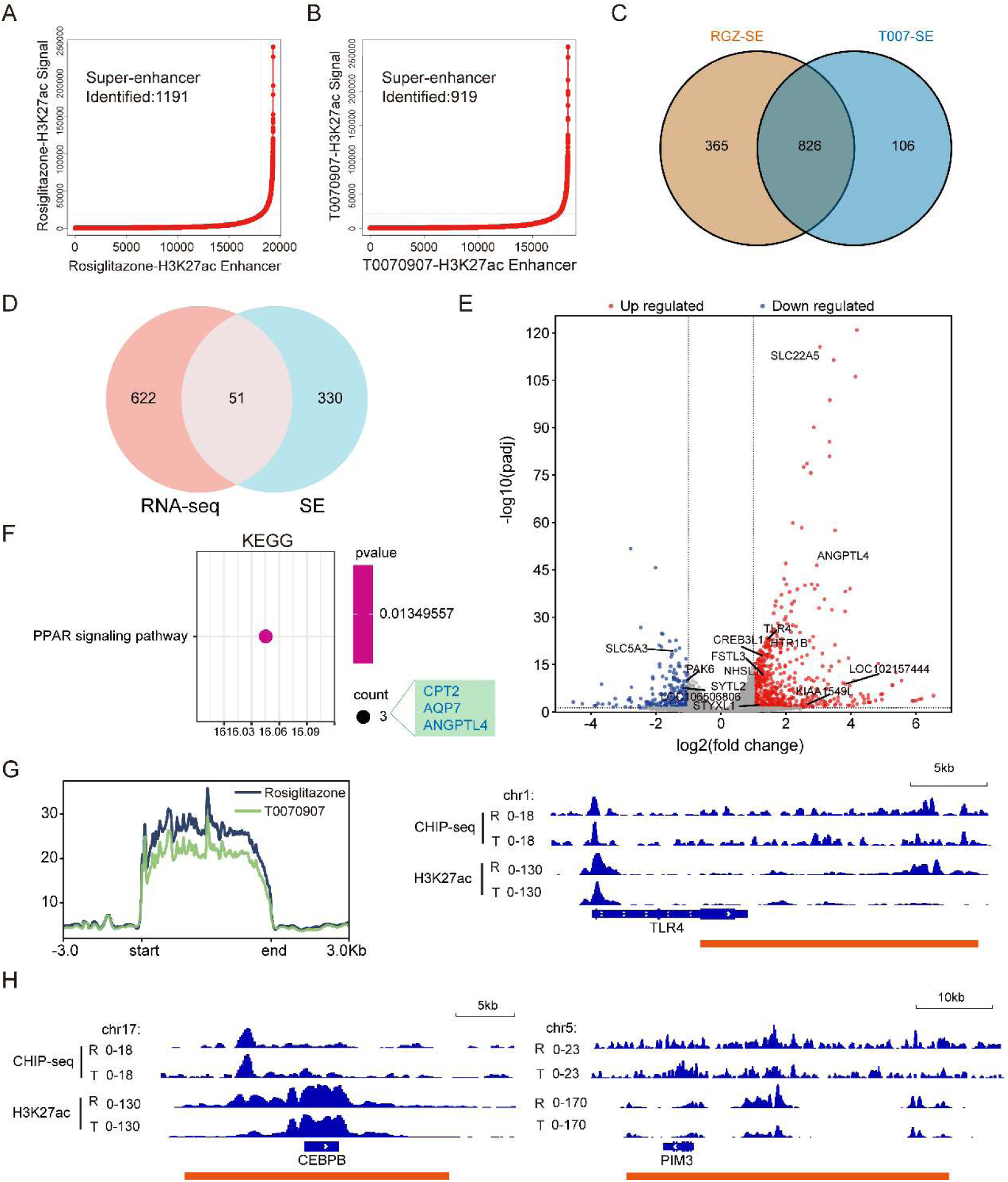
Activation of PPARγ remodels the super-enhancer landscape to regulate downstream target genes. **(A–B)** Identification of super-enhancers based on H3K27ac ChIP-seq signals in WSL-PPARG cells treated with RGZ (A) or T 007 (B). **(C)** Venn diagram of super-enhancer-associated genes under RGZ and T007 conditions, showing the numbers of treatment-specific and shared regulated genes. **(D)** Venn diagram showing the intersection analysis between RGZ-specific super-enhancer-associated genes and RNA-seq differentially expressed genes. **(E)** Visualization of RGZ-specific super-enhancers in the volcano plot of RNA-seq differentially expressed genes. **(F)** KEGG pathway enrichment analysis of the intersecting genes from (D). **(G)** Average ChIP-seq signal density of H3K27ac around the centers of super-enhancer regions. **(H)** IGV visualization of three RGZ-specific super-enhancers regulating PIM3, TLR4, and CEBPB. Super-enhancers were identified using the default inflection point of the ROSE algorithm. A corrected P-value (padjust) ≤ 0. 05 was considered statistically significant for enrichment analysis. ChIP-seq was performed with two biological replicates per group, and RNA-seq with three biological replicates per group, each with three technical replicates.

To elucidate the function of these RGZ-specific super enhancers, we integrated their associated genes with transcriptome data **(Fig. 7D-E)**. We identified 51 genes that were both differentially expressed and associated with RGZ-specific super-enhancers. Volcano plot analysis revealed that genes such as ANGPTL4 and SLC22A5 were significantly upregulated, while CREB3L3 and FSTL1 were downregulated. KEGG pathway enrichment analysis clearly indicated that these genes were most significantly enriched in the PPAR signaling pathway and included core lipid metabolism genes such as CPT2 and ANGPTL4 **(Fig. 7F)**. This demonstrates that PPARγ strongly drives its own metabolic reprogramming program by constructing super enhancers as master regulators.

The key finding is that this super-enhancer network deeply integrates core nodes of both innate immunity and inflammatory signaling, and analysis of H3K27ac signaling in super-enhancer regions shows that RGZ treatment significantly enhances epigenetic activation signals in these regions **(Fig. 7G)**, achieving a coupling of binding and activation states. IGV visualization further revealed that RGZ treatment induced significant H3K27ac super enhancer peaks near key gene loci of multiple immune signaling networks, such as the upstream pattern recognition receptor TLR4, the core inflammatory transcription factor CEBPB, and the signal regulatory kinase PIM3 **(Fig. 7H)**, which control initiation (TLR4), amplification (CEBPB), and regulation (PIM3) of inflammatory responses, respectively.

Based on these integrated analyses, we conclude that the core mechanism of PPARγ as a master regulator at the three-dimensional genome level: its activation can build super-enhancers on a large scale. This regulatory model, which closely couples metabolic programming with inflammation suppression through a shared super-enhancer architecture, provides a novel three-dimensional genomic explanation for how PPARγ integrally controls macrophage functional reprogramming.

## Discussion

This study reveals a core function of PPARγ in macrophages responding to cytoplasmic DNA (poly(dA:dT)) stimulation: it acts as a critical “brake” that integrates metabolic reprogramming with inflammation suppression through a multi-level, spatially structured genomic and epigenetic regulatory program. Specifically, activation of PPARγ achieves: (1) direct transcriptional repression of key genes in core inflammatory pathways (e.g., MAPK), such as DUSP4/10; (2) widespread reprogramming of the enhancer landscape by occupying distal regulatory elements and remodeling their H3K27ac modifications; and (3) driving the construction of novel super-enhancers that couple lipid metabolic pathways (e.g., CPT2) with core nodes of immune networks (TLR4, CEBPB, PIM3) within the same three-dimensional chromatin hub. This defines a systemic immune-metabolic integration mechanism mediated by PPARγ, spanning from one-dimensional sequence binding to 3D architectural reorganization.

PPARγ is a direct transcriptional regulator of inflammatory signaling(Costa et al, 2010). Traditional views emphasize PPARγ’s indirect anti-inflammatory effects via metabolic reprogramming or trans-repression with factors like NF-κB(Ricote & Glass, 2007). Our study updates this understanding through direct genomic evidence: activated PPARγ binds to promoters and enhancers of MAPK pathway negative regulators, cytoplasmic DNA sensors, and downstream inflammatory mediators via canonical peroxisome proliferator-activated receptor response elements (PPREs), enabling sequence-specific targeted transcriptional repression. Its ability to directly bind and downregulate genes such as the DUSP family indicates that PPARγ actively reprograms rather than passively antagonizes signaling networks, establishing rapid and sustained inhibitory states by upregulating endogenous negative feedback loops.

This direct regulation is implemented and reinforced through multi-level genomic events. PPARγ binding exhibits spatial diversity, occurring simultaneously at proximal promoter regions and distal/intergenic enhancer regions(Lefterova et al, 2014; Lefterova et al, 2008). This dual occupancy ensures comprehensive regulation from signal initiation to amplification. More importantly, its binding is tightly coupled with dynamic changes in local chromatin states—particularly at distal regulatory regions, where PPARγ recruitment is accompanied by downregulation (at inflammatory genes) or upregulation (at metabolic genes) of the H3K27ac active mark. This suggests PPARγ directly remodels the epigenetic landscape by recruiting co-regulatory complexes (e.g., corepressors or histone deacetylases), locking transcription factor binding into stable chromatin states (repressive or activating) to enable programmed co-expression of hundreds of genes(Hernandez-Quiles et al., 2021).

The most striking breakthrough of this study lies in revealing PPARγ-driven 3D genome reorganization, which couples immune and metabolic programs via super-enhancers. Super-enhancers are chromatin regulatory hubs formed by clusters of multiple typical enhancers, which are essential for maintaining cell identity and driving expression of core genes(Jiang et al., 2021). We found that PPARγ activation induces the formation of numerous de novo super-enhancers. These higher-order regulatory hubs are functionally versatile, elegantly integrating the drive of its own metabolic program (e.g., fatty acid oxidation genes) with targeting of multi-level components of immune signaling networks (upstream receptor TLR4, core transcription factor CEBPB, regulatory kinase PIM3). Among them, TLR4 serves as a critical upstream receptor that activates the downstream MAPK/NF-κB inflammatory signaling pathway(Nyati et al, 2017); CEBPB is a transcription factor that regulates the expression of various inflammatory cytokines, including TNF-α and IL-6(He et al, 2025; Wang et al, 2025) and PIM3 kinase has also been reported to be involved in cell survival and the regulation of inflammatory signaling(Baek et al, 2020; Mukaida et al, 2011). This provides the first molecular evidence for the immunometabolism concept at the 3D genome level in the context of cytoplasmic DNA responses. It clarifies that PPARγ not only influences metabolism and inflammation through linear pathways but also acts as a true integrator by constructing advanced chromatin architectures that enforce physical and functional synergy between metabolic reprogramming and inflammation suppression.

Our findings align with PPARγ’s known anti-inflammatory roles but greatly expand and deepen its mechanistic framework(Mukohda & Ozaki, 2021; Puhl et al, 2015). While confirming the efficacy of PPARγ agonists in suppressing inflammation, this study extends their functional context to the critical innate immune response of cytoplasmic DNA sensing. More importantly, it resolves the debate over whether its effects are “metabolically indirect” or “direct transcriptional”: our data show these are inseparable and unified via the super-enhancer architecture. Compared to previous models focusing on interactions with corepressors (e.g., NCoR)(Pascual et al, 2005), we reveal that direct DNA binding, concomitant epigenetic remodeling, and higher-order structural assembly are more fundamental mechanisms.

Elucidation of this multi-level regulatory mechanism holds significant biological and therapeutic implications. Physiologically, it explains how macrophages initiate potent antiviral (cytoplasmic DNA-sensing) responses while preventing excessive immune damage and apoptosis via a preset, multi-level braking program activated by PPARγ, thereby maintaining homeostasis(Yoon et al, 2015). Pathologically, it offers new insights into the “metabolism-inflammation” vicious cycle in chronic diseases like atherosclerosis and non-alcoholic fatty liver disease(Hulsmans & Holvoet, 2010): dysfunction of PPARγ may disrupt its integrated 3D regulatory network. Therapeutically, this study not only provides mechanistic support for existing PPARγ agonists (e.g., thiazolidinediones(Liu et al, 2020)) in treating diseases driven by cytoplasmic DNA accumulation (e.g., autoimmune diseases, certain DNA virus infections) but also points to novel precision interventions. For example, targeting PPARγ-regulated specific super-enhancers or key node genes (e.g., TLR4 enhancers) may facilitate the development of less toxic, more targeted immunometabolic modulators. This study has certain limitations and points to future directions. First, conclusions are primarily based on in vitro macrophage models; whether PPARγ’s regulatory network is conserved across tissue-resident macrophages or in complex in vivo inflammatory environments requires further validation. Second, we have not fully elucidated the specific cofactor complexes recruited by PPARγ or how they precisely regulate dynamic H3K27ac changes. Future studies should utilize CRISPR-based genome editing to validate the functional necessity of key super-enhancer elements in vitro and in vivo(Kravchuk et al, 2023); explore PPARγ’s 3D genome regulation in human disease-relevant cells (e.g., systemic lupus erythematosus)(Zhao et al, 2023); and investigate crosstalk between PPARγ and other nuclear receptors (e.g., LXR) in coordinating immunometabolic regulation.

In conclusion, this study reshapes the understanding of PPARγ as a classic nuclear receptor. We elevate it from a traditional metabolic regulator and indirect inflammation suppressor to an active “3D genome architect.” Through a programmed, spatially hierarchical multi-scale regulatory network—direct transcriptional repression, epigenetic landscape remodeling, and super-enhancer-mediated architectural coupling—it integrates macrophage metabolic states with inflammatory response intensity. This paradigm shift not only provides a novel framework for understanding immunometabolic crosstalk but also lays a solid theoretical foundation for developing next-generation therapeutics aimed at restoring this delicate balance(Wagner & Wagner, 2022).

## Materials and Methods

### Cell culture

The WSL-PPARG (iPAM) cell line was established in our laboratory(Wagner & Wagner, 2022). Cells were cultured in DMEM (Corning, 10-013-CVRC) and RPMI 1640 (HyClone, SH30809.01) mixed at a 1:1 ratio, supplemented with 10% fetal bovine serum and 1% penicillin-streptomycin. Cultures were maintained at 37℃ in a humidified incubator with 5% CO2.

### Effects of PPARG activation and inhibition on poly(dA:dT) transfection

iPAM cells were seeded and pre-treated with DMSO (Solarbio, D8371), Rosiglitazone (Sigma, R2408), or T0070907 (Selleck Chemicals, S2871) for 4 h. When cells reached 80%-85% confluence, transfection was performed using Lipofectamine 2000 (Thermo Fisher Scientific, 11668019). For a 60 mm dish, 30.5 µL of Lipofectamine 2000 and 4 µg of Poly(dA:dT) naked ‘InvivoGen, tlrl-patn’ were each diluted in 366 µL of Opti-MEM, gently mixed, and incubated at room temperature for 5 minutes to form the transfection complexes. The complexes were then added dropwise to the cell culture medium, and cells were cultured for an additional 4 h before sample collection for analysis.

### Detection of poly(dA:dT)-induced apoptosis by hoechst-PI fluorescence staining

iPAM cells were seeded in 6-well plates and pre-treated with DMSO, Rosiglitazone, or T0070907 for 4 h. Poly(dA:dT) naked was transfected into the cells using Lipofectamine 2000. At 24 h post-transfection, apoptosis was assessed by fluorescence microscopy and flow cytometry. For fluorescence microscopy, cells were washed three times with PBS (pH 7.4), stained with 5 μL of Hoechst staining solution and 5 μL of PI staining solution (Beyotime, C1056) at 4℃ for 30 min, washed once with PBS, and then observed. For flow cytometry, adherent cells were collected, resuspended in 1 mL of cell staining buffer (Beyotime, C1056), stained with the same dye solutions as described above, incubated, and directly analyzed by flow cytometry. Both methods utilized Hoechst-PI double staining to evaluate the morphological changes associated with apoptosis and necrosis induced by Poly(dA:dT).

### RNA-Seq analysis

For transcriptome analysis, iPAM cells were pre-treated with DMSO, Rosiglitazone (10 μM), or T0070907 (2 μM) for 4 h, followed by transfection with Poly(dA:dT) naked using Lipofectamine 2000. Total RNA was extracted from iPAM cells using TRIzol reagent, and RNA integrity was assessed using the Bioanalyzer 2100 system (Agilent Technologies, Santa Clara, CA, USA). All RNA samples were subsequently sent to Novogene (Beijing, China) for transcriptome sequencing.

Raw sequencing data (raw reads) were initially processed using fastp software to remove adapter-containing reads, poly-N reads, and low-quality reads, yielding clean reads. The pig (Sus scrofa) genome (assembly Sscrofa11.1) index was built using Hisat2 v2.0.5, and paired-end clean reads were aligned to the reference genome using the same software. Gene expression levels were quantified as fragments per kilobase of transcript per million mapped reads (FPKMs) using FeatureCounts v1.5.0-p3. Differential expression analysis among the four experimental groups was performed using the DESeq2 R package (version 1.20.0).

### ChIP-seq analysis

For chromatin immunoprecipitation followed by sequencing (ChIP-seq), cells at 90%-95% confluence were cross-linked with 1% formaldehyde at room temperature for 10 min, and the reaction was terminated with 125 mM glycine. Cells were washed with ice-cold PBS, scraped, collected, and stored. Subsequently, cells were lysed and sonicated to fragment DNA into 200 – 1000 bp fragments. Target protein-DNA complexes were immunoprecipitated using specific antibodies, followed by stringent washing and elution. The recovered complexes were de-crosslinked, and DNA was purified. Purified DNA fragments were end-repaired, A-tailed, and ligated to sequencing adapters. Libraries were constructed by agarose gel purification and PCR amplification. After quality assessment, libraries were sequenced on an Illumina platform. Raw sequencing data were quality-controlled using fastp, and clean reads were aligned to the reference genome (Sscrofa11.1). The distribution of mapped reads across genomic regions, including coding sequences (CDS), intergenic regions, and introns, was analyzed to determine their origin. Peak calling was performed using MACS2 (for transcription factors) or Epic2 (for histones) with input samples as controls. Finally, Gene Ontology (GO) and Kyoto Encyclopedia of Genes and Genomes (KEGG) pathway enrichment analyses of peak-associated genes were performed and visualized using an online data analysis and visualization platform (https://www.bioinformatics.com.cn). Gene peak visualization was conducted using Integrative Genomics Viewer (IGV) software.

### Identification of SEs

To identify super-enhancers in porcine alveolar macrophages under conditions of PPARG activation and inhibition, we conducted an analysis following established methodologies from published research. Initially, super-enhancers were identified in raw form using ROSE software with the parameters -s 12500 -t 3000, applied to H3K27ac ChIP-seq data from porcine alveolar macrophages treated with T0070907, Rosiglitazone, and their corresponding control groups. Specifically, the normalized read counts per million mapped reads were calculated for each enhancer region and adjusted based on enhancer length. Enhancers located above the inflection point were classified as super-enhancers. Finally, the genes nearest to the super-enhancers identified by ROSE were selected as the target genes regulated by these super-enhancers.

### Statistical analysis

All experiments were independently repeated at least three times. Data are expressed as mean ± SD and analyzed using GraphPad Prism 9.0. Statistical comparisons between two groups were performed with two-tailed Student’s t-test, while multiple-group comparisons were evaluated by one-way or two-way ANOVA as appropriate. Significance levels are indicated as ns (not significant), \**p* ≤ 0.05, \*\**p* ≤ 0.01, \*\*\**p* ≤ 0.001, and \*\*\*\**p* ≤ 0.0001.

## Supplementary materials

Fig. S1: Genomics of PPARγ regulating the PPAR signaling pathway in non-coding regions.

## Data availability

The RNA-seq and ChIP-seq data generated in this study have been deposited in the Genome Sequence Archive (GSA) of the National Genomics Data Center, China National Center for Bioinformation, under the accession number (CRA025774). All other data supporting the findings of this study are available from the corresponding author upon reasonable request.

## Author contributions

**Xinyi Fan**: Formal analysis; Investigation; Methodology; Writing— original draft; Writing—review and editing. **Kai Gao**: Software; Formal analysis; Writing—review and editing. **Ping Meng**: Formal analysis; Investigation; Writing—review and editing. **Jingmei Xu**: Investigation; Data curation. **Gongrui Zhang**: Resources; Investigation. **Yumei Yuan**: Investigation. **Qifeng Gao**: Software; Formal analysis; Investigation. **Zhuoyuan Ma**: Data curation; Software. **Fan Yang**: Resources. **Yongqiang Zhang**: Supervision; Funding acquisition; Investigation; Project administration. **Yan Wang**: Conceptualization; Formal analysis; Supervision; Writing—original draft. **Shiqiang Zhang**: Conceptualization; Project administration; Funding acquisition; Supervision; Writing—review and editing.

## Acknowledgements

We are grateful to the members of the Shaanxi Engineering Research Center of Stem Cell for sharing valuable materials and their technical support. We gratefully acknowledge the National Natural Science Foundation of China (Grant No. 32072856) support for this research.

---

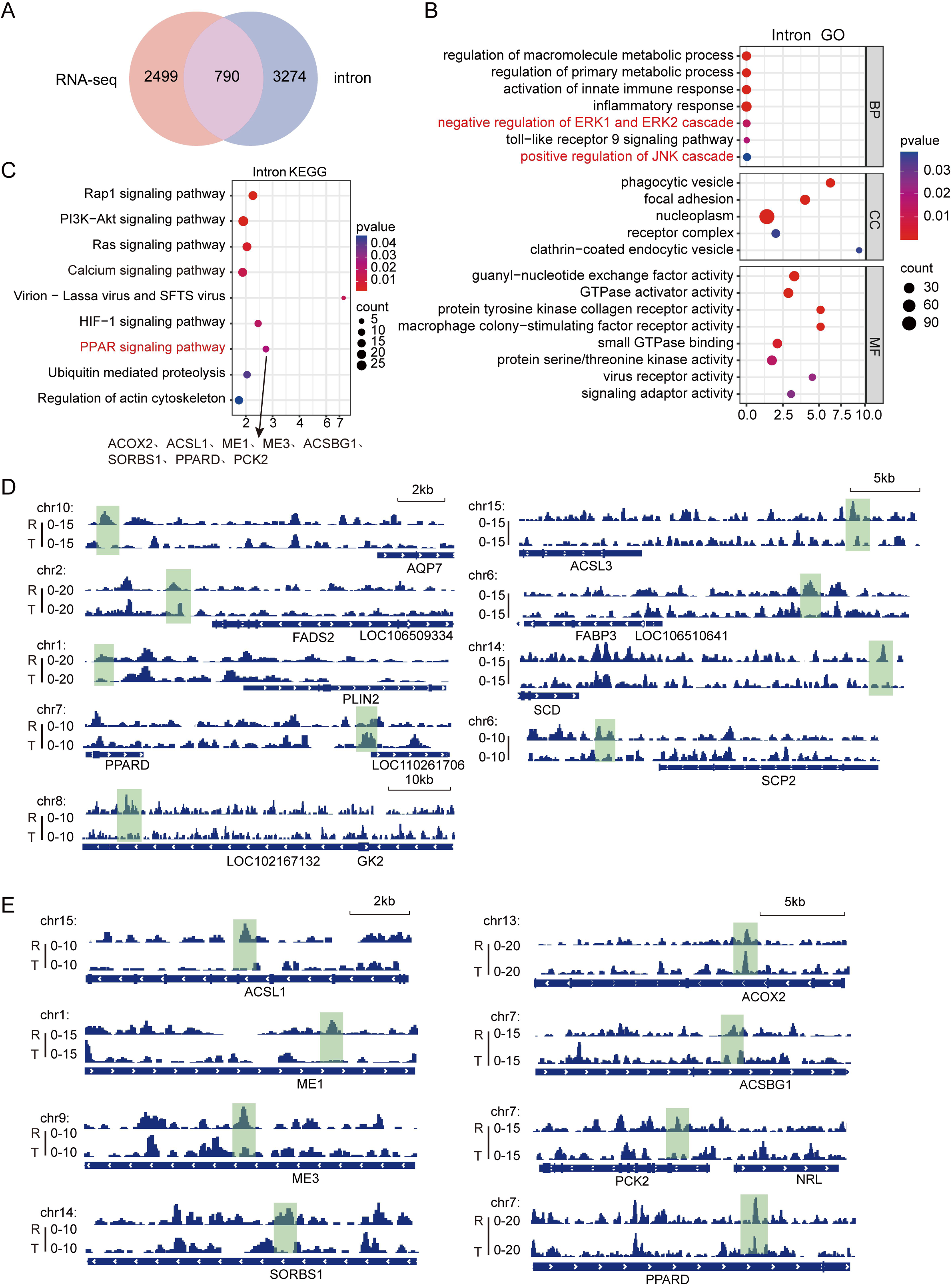

